# Anti-V2 Antibody Deficiency in Individuals Infected With HIV-1

**DOI:** 10.1101/336339

**Authors:** Lily Liu, Liuzhe Li, Aubin Nanfack, Luzia M. Mayr, Sonal Soni, Adam Kohutnicki, Lucy Agyingi, Xiao-Hong Wang, Michael Tuen, Yongzhao Shao, Maxim Totrov, Susan Zolla-Pazner, Xian-Peng Kong, Ralf Duerr, Miroslaw K. Gorny

## Abstract

The positive correlation of high levels of plasma anti-V2 antibodies (Abs) with protective immunity in the Phase III anti-HIV RV144 vaccine trial generated interest in the induction of these Abs for HIV vaccine development. We analyzed plasma samples from 79 chronically infected Cameroonian individuals for Ab reactivity against three V1V2 fusion proteins and five cyclic V2 peptides and found that HIV-1 infection induces different levels of anti-V2 Abs. While the majority of plasma samples reacted strongly with one or more V2 antigens, 10% (8) of the samples were nonreactive. Deficiency of anti-V2 Abs was consistently found in longitudinal plasma samples tested over 8 to 54 months of HIV infection. There was a strong correlation between binding activities of plasma anti-V2 Abs and anti-gp120 and anti-gp41 Abs, suggesting that deficiency of V2 Abs could be related, in part, to a limited ability to elicit strong Ab responses. Analysis of gp120 sequences revealed that the V2 region of viruses from donors with V2-deficient versus V2-reactive Abs displayed a tendency toward longer length, more glycans, and lower isoelectric point and charge. No differences between these two patient groups were noted in the same parameters measured in the V1 region. These data suggest that immunogens containing a shorter V2 region with fewer glycosylation sites and higher electrostatic charges would be beneficial for induction of anti-V2 Abs, but the ability to mount a strong general Ab response to HIV-1 appears to be a dominant factor.

**IMPORTANCE:** The results of the RV144 vaccine clinical trial showed a correlation between plasma Abs against a V1V2 fusion protein and a decreased risk of acquiring HIV-1 infection. This turned the focus of some HIV vaccine design to the induction of elevated levels of anti-V2 Abs to increase vaccine efficacy. In plasma samples from Cameroonian individuals infected with HIV-1, we observed broad variations in levels of anti-V2 Abs, and 8 of the 79 plasma samples tested displayed substantial deficiency of V2 Abs. Sequence analysis of the V2 region from plasma viruses and multivariate analyses of V2 characteristics showed a significant difference in several features between V2-deficient and V2-reactive plasma Abs. These results suggest that HIV vaccine immunogens containing a V2 region with shorter length, fewer glycosylation sites, and higher electrostatic charges may be beneficial for induction of a higher level of anti-V2 Abs and thus contribute to HIV vaccine efficacy.

---

Anti-V2 antibodies (Abs) have been the subject of studies to determine their role in protection against HIV-1 infection due to the results of the Phase III RV144 vaccine trial, which showed an inverse correlation with risk of infection (1). The RV144 data revealed that the V2 Abs in vaccinees were highly cross-clade reactive and that high levels, (but not medium or low levels) of plasma Abs against a V1V2 fusion protein correlated with reduced acquisition of HIV-1 infection (1, 2). The V2 region contains at least 3 types of epitopes defined by anti-V2 monoclonal Abs (mAbs): i) V2 integrin (V2i), which is a conformation-dependent motif and includes the α4β7 integrin binding site; ii) the V2 peptide (V2p) linear epitope; and iii) the V2 quaternary (V2q) epitope, present preferentially on the trimer (3). The majority of anti-V2 Abs are specific for conformation-dependent V2i epitopes; a smaller amount to the linear V2p epitopes, and some are specific for quaternary V2q epitopes and only present in a small proportion of HIV-infected individuals.

The frequency of anti-V2 Abs in the plasma of individuals infected with HIV-1 varies among studies, depending on the methods used and the population studied. Screening of immune plasma using V2 consensus peptides with the HxB2 sequence detected 12% and 21% positive sera from HIV-1-infected individuals (4, 5). Using V1V2 proteins expressing both conformation-dependent V2i and linear V2p epitopes, the frequency of plasma anti-V2 Abs is higher, yielding 30% and 48% in two studies (6, 7). The prevalence of anti-V2 Abs in clade B-infected individuals was determined to be 45% when tested against a C1-V1V2 BH10 protein and 30% when screened in a competition assay with the 697 mAb specific for the conformational V2i epitope (8).

The frequency of anti-V2 Abs induced by the RV144 vaccine was relatively high, given that 84% of vaccinees’ sera was reactive with the V1V2-gp70 fusion protein carrying the sequence of subtype B strain CaseA2 (9). A high percentage of plasma samples was also reactive with the cyclic V2 peptide with the sequence of clade AE 92TH023 (1). Given that the high level of anti-V2 Abs is relevant for protection (2) and that not all immune plasma samples from infected and vaccinated individuals contain V2 Abs, we studied a panel of plasma samples to determine the frequency of anti-V2 Abs against different V1V2 fusion proteins and cyclic V2 peptides. Lack of anti-V2 Abs in some plasma samples was in part dependent on a weak general Ab response to HIV envelope (Env) antigens, manifested by significantly lower titers of Abs against V3, gp120, and gp41 proteins compared with plasma with reactive anti-V2 Abs. In addition, differences in the V2 region, including extended length, glycosylation sites, lower isoelectric point, and charge may also contribute to lower immunogenicity and deficiency of anti-V2 Abs in some individuals infected with HIV-1.

## RESULTS

### Frequency of plasma Abs against V2 and control antigens

A panel of 79 plasma samples was obtained at the Medical Diagnostic Center in Yaoundé, Cameroon from a cohort of individuals chronically infected with HIV-1. The plasma samples were screened at a 1:100 dilution by ELISA against three V1V2 fusion proteins and five biotinylated cyclic V2 peptides. Given that anti-V1 Abs are sequence-specific (10) and don’t bind to heterologous V1V2 fusion proteins, plasma samples can be tested against V1V2 fusion proteins with heterologous sequence to detect specific Abs against the V2 region. To confirm that the anti-V1 Abs would not bind to heterologous V1V2 fusion proteins, we screened all plasma samples against two fusion proteins with the major parts of V2 deleted: V1ΔV2_A244_-gp70 and V1ΔV2_CaseA2_-gp70 (Table S1). None of the 79 plasma samples reacted with these two V1ΔV2 proteins, confirming that binding of plasma to heterologous V1V2 fusion proteins usually detects Abs specific for V2.

The frequency of plasma with anti-V2 Abs varied depending on the antigen sequence. The highest percentage of plasma samples reacted with V1V2_case A2_-gp70 at 85%; 80% reacted with V1V2_A244_-gp70; and 53% of plasma Abs bound to V1V2_ZM109_-1FD6 fusion proteins (Fig. 1A). The frequency of anti-V2 peptide Abs was lower, ranging from 72% to 40% for plasma binding to V2_A244_, V2_92TH023_ (clade AE), V2_Du422_ (clade C), V2_230_ and V2_200_ (clade AG) (Fig. 1A).

**Figure 1.**
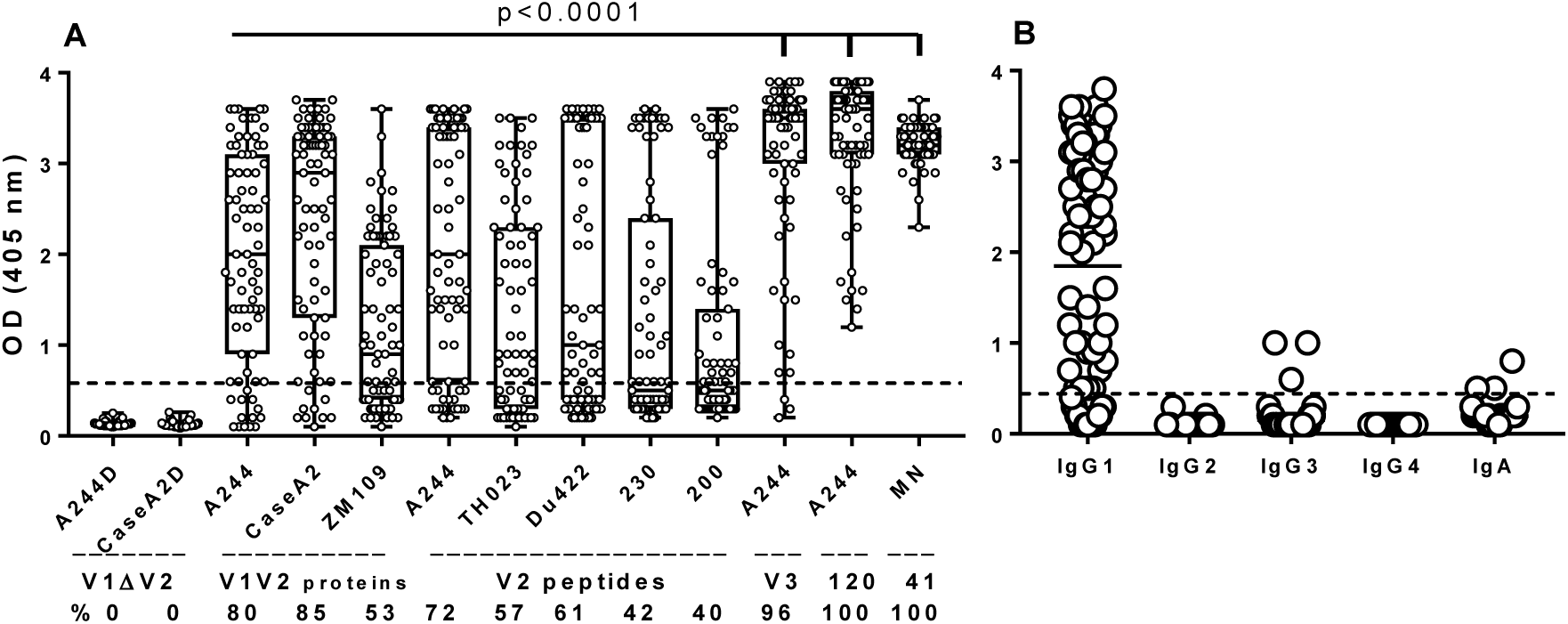
Reactivity of plasma Abs against V2 antigens and control proteins. (A) All 79 plasma samples from Cameroonian HIV-1 infected individuals were tested at 1:100 dilutions by ELISA against proteins coated at 1 μg/mL. Antigens included: two V1V2-gp70 fusion proteins with V2 deleted (V1ΔV2) with sequences from A244 (clade AE) and CaseA2 (clade B); three V1V2 fusion proteins: V1V2_A244_-gp70 (clade AE), V1V2_Case A2_-gp70 (clade B) and V1V2_ZM109_-1FD6 (clade C); five biotinylated cyclic V2 peptides with sequences from A244 (clade AE), TH023 (clade AE), Du422 (clade C), 230 and 200 (clade AG); one biotinylated cyclic V3_A244_ peptide, gp120_A244_ protein and gp41_MN_ protein. Percentage of plasma samples with specific anti-V2 Abs is shown below each antigen. Statistical significance between Abs against V2 and control antigens was determined by nonparametric Mann-Whitney test. The shape of the distribution is shown by a box with the ends of the box representing the 5th and 95th percentile, and the median marked by a horizontal line inside the box. (B) IgG subclasses and IgA of plasma Abs binding to V1V2_A244_-gp70 fusion protein.

All 79 plasma samples reacted by ELISA with gp120_A244_ and gp41_MN_ proteins, while 76 (96%) bound to cyclic V3_A244_ peptide, most of the latter reaching the saturation level at OD 3.5 to 3.9 (Fig. 1A). The IgG subclasses and IgA of plasma Abs, tested separately against V1V2_A244_-gp70, showed the dominance of IgG1 Abs, whereas IgG3 and IgA were only detected in 3 plasma samples each, and IgG2 and IgG4 binding was not detectable (Fig. 1B). The results showed a broad range of ODs for anti-V1V2 fusion proteins and anti-V2 peptides Abs, from close to 0 to 3.9, while the OD range for anti-gp120 was from 1.2 to 3.9 and for anti-gp41 from 2.1 to 3.7 OD (all positive). This indicates a weaker immunogenicity of the V2 region compared with gp120 and gp41, which was further confirmed by 50% titers of plasma Abs, as shown below.

The CD4^+^ T cell counts, cells/μL, were comparable in two tested groups (6 samples each): V2 deficient and V2 reactive; mean counts were higher but not significant in the V2-deficient group (558 vs 402, respectively; *P*=0.1426). Viral loads, HIV RNA copies/mL, were measured in the same plasma samples, and results in the V2-deficient and V2-reactive groups were also comparable: 7,076 vs 16,400, respectively (*P*=0.2780).

Glycoprotein 120 from 12 plasma viruses (6 each from the V2-deficient and V2-reactive groups) was sequenced, and virus subtypes were identified (Table 1). All but one virus (clade C) were HIV-1 circulating recombinant forms (CRF), which dominate in Cameroon, according to recent analysis with the most prevalent CRF02_AG (64.9%) and CRF22_01A1 (7.1%) (11, 12). The identified subtypes in this study were as follows: CRF02_AG (3 plasma samples), CRF01_AE (2), CRF22_01A1 (2), CRF11_cpx, CRF18_cpx, CRF36_cpx, and CRF37_cpx (Table 1).

**TABLE 1.**
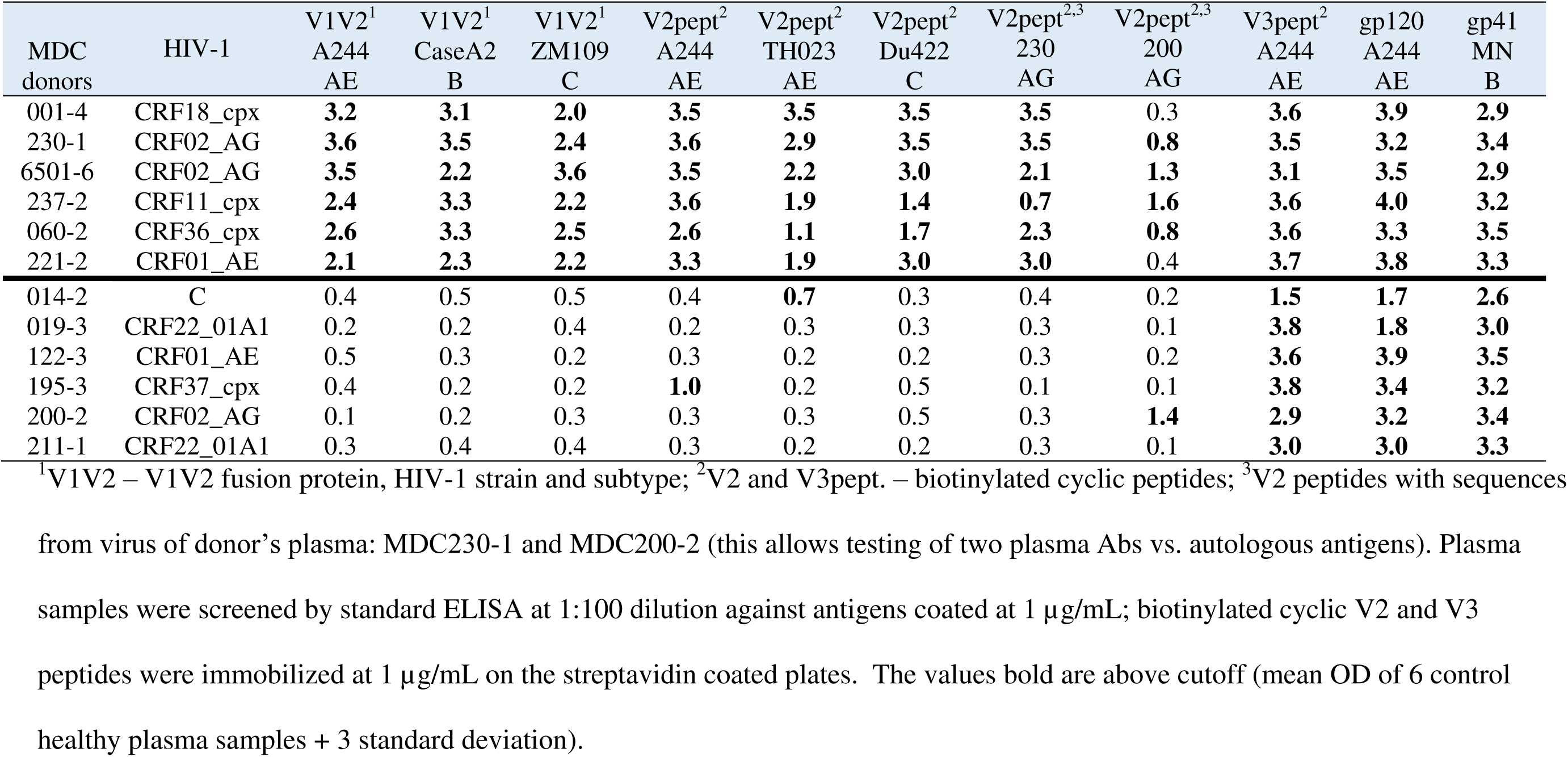
Infecting HIV clades and ELISA reactivity of plasma antibodies to Env antigens of patients with and without V2 antibodies

| MDC donors | HIV-1 | V1V2<sup>1</sup><br>A244<br>AE | V1V2<sup>1</sup><br>CaseA2<br>B | V1V2<sup>1</sup><br>ZM109<br>C | V2pept<sup>2</sup><br>A244<br>AE | V2pept<sup>2</sup><br>TH023<br>AE | V2pept<sup>2</sup><br>Du422<br>C | V2pept<sup>2,3</sup><br>230<br>AG | V2pept<sup>2,3</sup><br>200<br>AG | V3pept<sup>2</sup><br>A244<br>AE | gp120<br>A244<br>AE | gp41<br>MN<br>B |
| --- | --- | --- | --- | --- | --- | --- | --- | --- | --- | --- | --- | --- |
| 001-4 | CRF18_cpx | **3.2** | **3.1** | **2.0** | **3.5** | **3.5** | **3.5** | **3.5** | 0.3 | **3.6** | **3.9** | **2.9** |
| 230-1 | CRF02_AG | **3.6** | **3.5** | **2.4** | **3.6** | **2.9** | **3.5** | **3.5** | **0.8** | **3.5** | **3.2** | **3.4** |
| 6501-6 | CRF02_AG | **3.5** | **2.2** | **3.6** | **3.5** | **2.2** | **3.0** | **2.1** | **1.3** | **3.1** | **3.5** | **2.9** |
| 237-2 | CRF11_cpx | **2.4** | **3.3** | **2.2** | **3.6** | **1.9** | **1.4** | **0.7** | **1.6** | **3.6** | **4.0** | **3.2** |
| 060-2 | CRF36_cpx | **2.6** | **3.3** | **2.5** | **2.6** | **1.1** | **1.7** | **2.3** | **0.8** | **3.6** | **3.3** | **3.5** |
| 221-2 | CRF01_AE | **2.1** | **2.3** | **2.2** | **3.3** | **1.9** | **3.0** | **3.0** | 0.4 | **3.7** | **3.8** | **3.3** |
| 014-2 | C | 0.4 | 0.5 | 0.5 | 0.4 | **0.7** | 0.3 | 0.4 | 0.2 | **1.5** | **1.7** | **2.6** |
| 019-3 | CRF22_01A1 | 0.2 | 0.2 | 0.4 | 0.2 | 0.3 | 0.3 | 0.3 | 0.1 | **3.8** | **1.8** | **3.0** |
| 122-3 | CRF01_AE | 0.5 | 0.3 | 0.2 | 0.3 | 0.2 | 0.2 | 0.3 | 0.2 | **3.6** | **3.9** | **3.5** |
| 195-3 | CRF37_cpx | 0.4 | 0.2 | 0.2 | **1.0** | 0.2 | 0.5 | 0.1 | 0.1 | **3.8** | **3.4** | **3.2** |
| 200-2 | CRF02_AG | 0.1 | 0.2 | 0.3 | 0.3 | 0.3 | 0.5 | 0.3 | **1.4** | **2.9** | **3.2** | **3.4** |
| 211-1 | CRF22_01A1 | 0.3 | 0.4 | 0.4 | 0.3 | 0.2 | 0.2 | 0.3 | 0.1 | **3.0** | **3.0** | **3.3** |

<sup>1</sup>V1V2 – V1V2 fusion protein, HIV-1 strain and subtype; <sup>2</sup>V2 and V3pept. – biotinylated cyclic peptides; <sup>3</sup>V2 peptides with sequences
from virus of donor's plasma: MDC230-1 and MDC200-2 (this allows testing of two plasma Abs vs. autologous antigens). Plasma samples were screened by standard ELISA at 1:100 dilution against antigens coated at 1 µg/mL; biotinylated cyclic V2 and V3 peptides were immobilized at 1 µg/mL on the streptavidin coated plates. The values bold are above cutoff (mean OD of 6 control healthy plasma samples + 3 standard deviation).

### Reactivity of plasma anti-V2 Abs

The majority of 79 plasma samples at 1:100 dilutions (90%) were cross-reactive against eight V2 antigens including three V1V2 fusion proteins and five V2 peptides and representing sequences from HIV-1 subtypes AE, AG, B and C. Eleven plasma samples (14%) reacted with all eight V2 antigens (Fig. 2A), while 60 plasma samples (76%) reacted with several, but not all, V2 antigens (data not shown). Eight plasma samples (10%) did not react with any of the eight V2 antigens (Fig. 2B) with the exception of three plasma samples which each bound weakly to one V2 antigen, but still reacted with cyclic V3_A244_ peptide, gp120_A244_ and gp41_MN_ (Fig. 1A).

**Figure 2.**
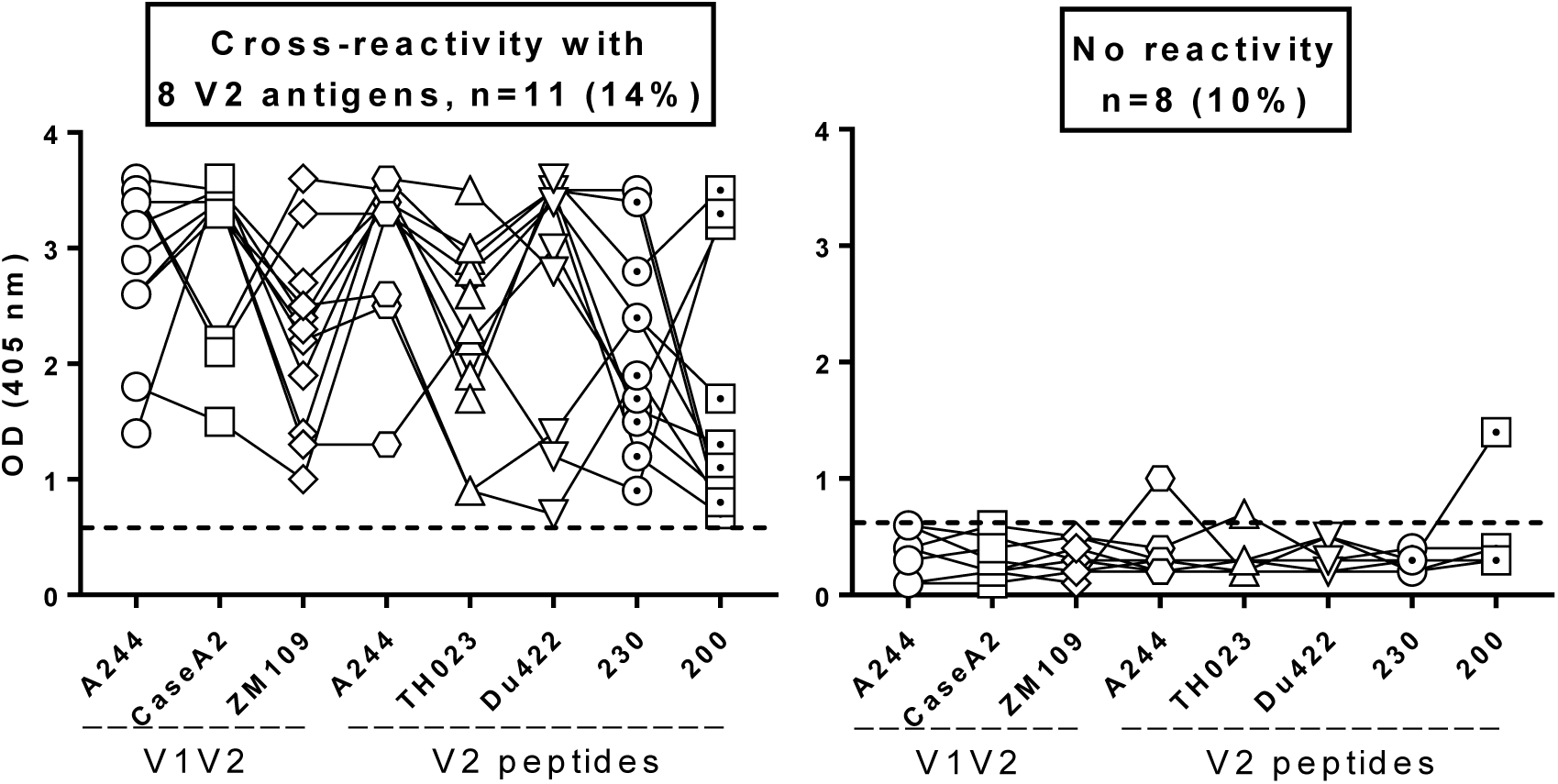
Cross-reactivity of plasma Abs with eight V2 antigens. (A) Eleven Cameroonian plasma samples (out of 79) at dilution 1:100 tested by ELISA demonstrate binding to all eight V2 antigens: V1V2A244-gp70 (clade AE), V1V2_CaseA2_-gp70 (clade B), V1V2_ZM109_-1FD6 (clade C), and biotinylated cyclic V2_A244_, V2_TH023_, (clade AE), V2_Du422_ (clade C), V2_230_ and V2_200_ (clade AG) peptides. (B) Eight plasma samples did not bind to V2 antigens tested with the exception of three plasma samples which each bound weakly to one V2 peptide. All plasma samples displayed reactivity with gp120_A244_ and gp41_MN_ (Fig.1). The dashed lines indicate the cutoff (mean OD + 3xSD of normal human plasma).

Given that plasma sample reactivity is usually tested against heterologous V2 sequences, we also included cyclic V2 peptides with sequences derived from two blood donors, MDC230 and MDC200, to test the reactivity to autologous linear V2 epitopes (Table 1). The results showed strong binding of plasma MDC230-1 (V2-reactive group) to the autologous V2_230_ peptide while plasma MDC200-2 (V2-deficient group) bound only weakly to the autologous V2_200_ peptide but not to other heterologous V2 antigens (Table 1). Of interest, the V2_200_ peptide was weakly reactive with 4 of 6 plasma samples from the cross-reactive group (Table 1). One unusual feature of the V2 region represented by the V2_200_ peptide is its exceptionally low pI of 4.79, while other V2 regions have pI values between 6.52 and 9.69 (Table S3).

### Longitudinal studies of plasma anti-V2 Abs

To determine whether the deficiency of V2 Abs is temporary or stable, we analyzed plasma samples from six V2 Ab-deficient donors over 11-54 months of observation and plasma samples from six V2 Ab-reactive donors over 28-60 months of observation (Table 1). These sequential samples were titrated and screened against five Env antigens: V1V2_A244_-gp70, cyclic V3_A244_ peptide, gp120_A244_, gp41_MN_, and p24_HXB2_ (Fig. 3). Titers of anti-V2_A244_ Abs in the V2-deficient group were undetectable (<1:100 dilution) (Fig. 3A) and designated as a 50% binding titer of 10^0^ (Fig. 4). Longitudinal analyses confirmed that lack of V2 Abs in plasma collected from six selected individuals infected with HIV-1 was constant over 11 to 54 months of observation. Similarly, Abs to four other antigens (V3, gp120, gp41, p24) were detected at relatively constant levels, albeit at intermediate to high titers; these results show an exclusive, constant lack of Ab response to V2 (Fig. 3A). The panel of sequential plasma samples collected over 28 to 60 months from six volunteers with reactive V2 Abs displayed relatively constant levels of all five types of Abs, including anti-V2 Abs (Fig. 3B).

**Figure 3.**
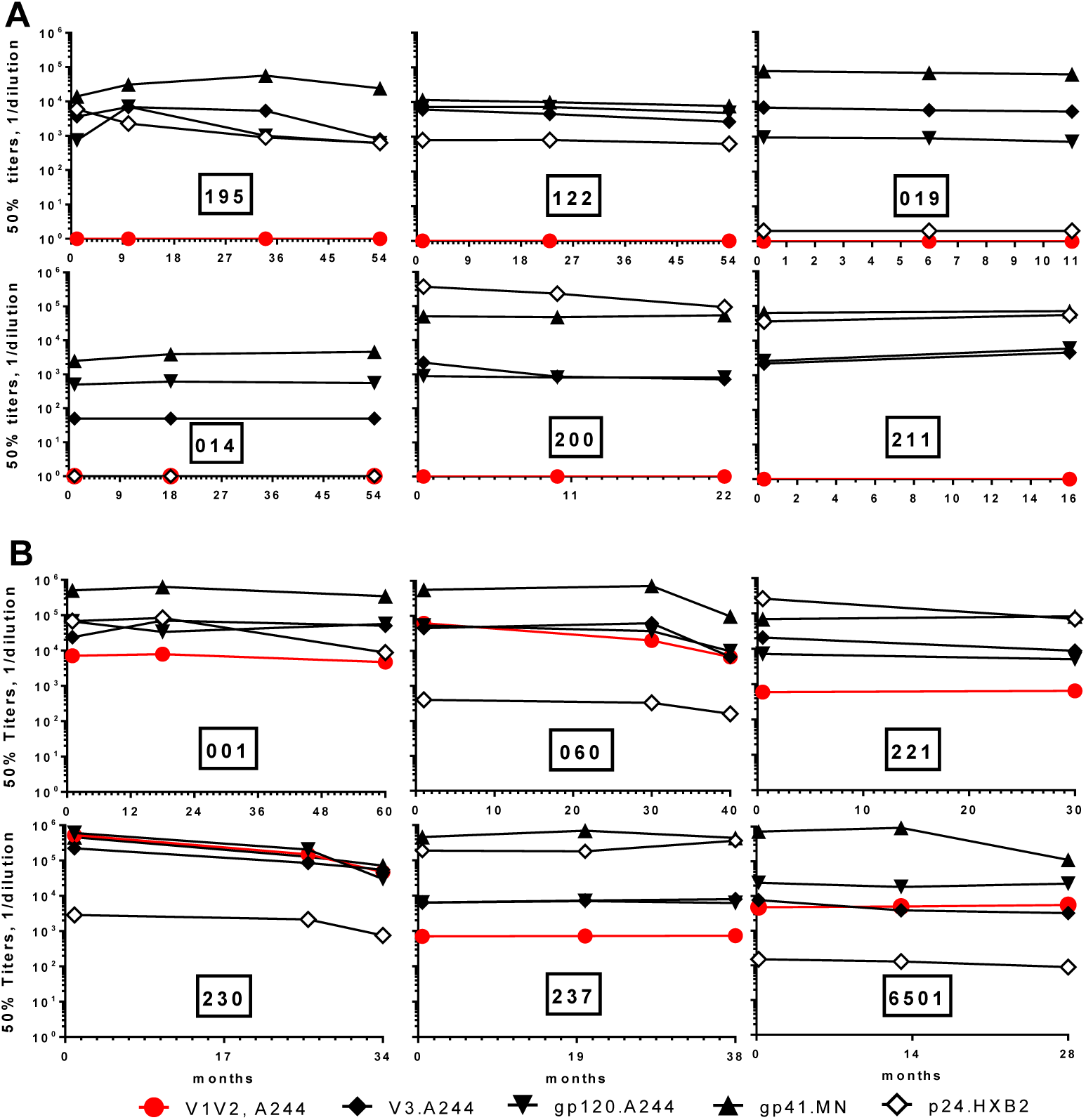
Titers of anti-V2 and control Abs in sequential plasma samples. (A) Six donors with lack of anti-V2 Abs over 11 to 54 months of observation, and (B) six donors with cross-reactive anti-V2 Abs over 28 to 60 months. The 50% titers of plasma Abs were determined by ELISA against V1V2_A244_-gp70 (clade AE) protein, biotinylated cyclic V3_A244_ peptide, gp120_A244_, gp41_MN_ (clade B), and p24_HXB2_ (clade B) protein. The antigens were coated or immobilized onto plate at 1 μg/mL; plasma were diluted by 10-fold dilutions starting from 1:100 to 1:1,000,000.

**Figure 4.**
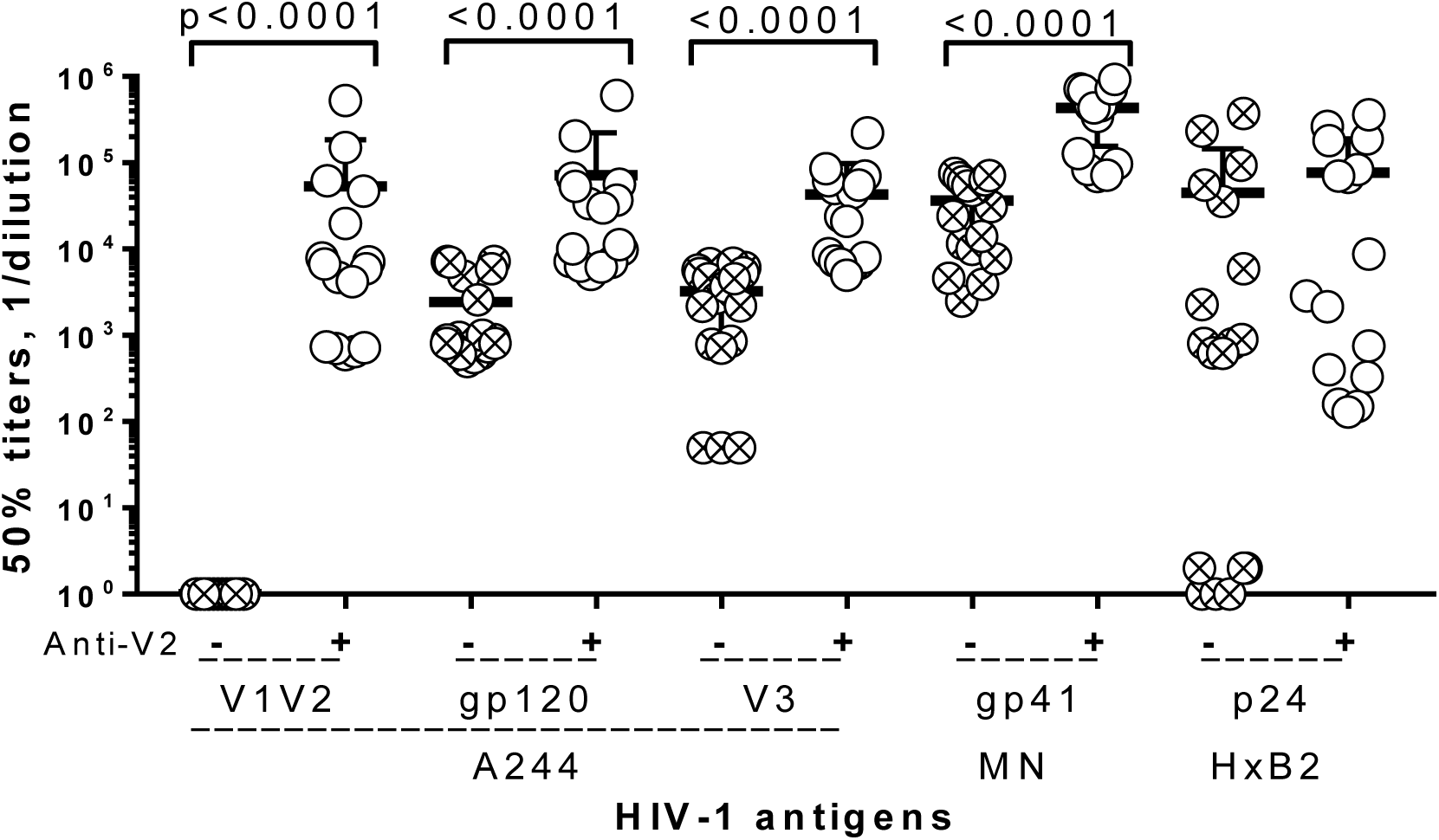
Titers of plasma Abs from HIV-1 infected individuals against HIV-1 antigens, V1V2, V3, gp120, gp41 and p24; two groups of plasma were compared: with non-V2-reactive (crossed circles) and cross-reactive anti-V2 Abs (open circles). The 50% titers of plasma Abs were determined by ELISA at five serial 10-fold dilutions starting from 1:100. Statistical significance was determined by nonparametric Mann-Whitney test.

The 50% titers of plasma Abs were used to compare the two groups of plasma samples (Fig. 4). There were significantly higher 50% titers of Abs (*P*<0.0001) against V1V2_A244_-gp70, cyclic V3_A244_ peptide, gp120_A244_, and gp41_MN_ in the group of V2-reactive Abs vs those with V2-deficitent Abs (Fig. 4, Table 2). Titers of anti-p24 Abs in the two groups were comparable (Fig. 4, Table 2).

**TABLE 2.** Titers of plasma Abs, CD4^+^ T cells, and viral load in two groups of HIV-1 infected subjects with anti-V2 deficient and cross-reactive Abs

| Features | V2 deficient<sup>1</sup><br>n=6 | V2 cross-reactive<sup>2</sup><br>n=6 | P value<sup>3</sup> |
| --- | --- | --- | --- |
| **50 % Titer<sup>4</sup>** |  |  |  |
| Anti-V1V2<sub>A244</sub> | 0.0 | 50,270 | **<0.0001** |
| Anti-V3<sub>A244</sub> | 3,252 | 40,572 | **<0.0001** |
| Anti-gp120<sub>A244</sub> | 2,438 | 70,241 | **<0.0001** |
| Anti-gp41<sub>MN</sub> | 36,444 | 413,169 | **<0.0001** |
| Anti-p24<sub>HXB2</sub> | 44,696 | 72,451 | 0.0781 |
| **CD4<sup>+</sup> T cells, VL** |  |  |  |
| CD4<sup>+</sup> T cell count | 558 | 402 | 0.1426 |
| Viral load (VL) | 7,076 | 16,400 | 0.2780 |

<sup>1</sup>Panel of plasma with undetectable anti-V2 Abs; <sup>2</sup>Panel of plasma with cross-reactive anti-V2 Abs; <sup>3</sup>nonparametric Mann-Whitney test; <sup>4</sup>mean of 50% titers.

### Correlation between Ab binding to V2 and other Env antigens

To determine if the deficit of anti-V2 Abs is related to the general Ab response to HIV-1 Env proteins, we analyzed the relationships of the ELISA binding activities (OD) between plasma Abs against V1V2_A244_-gp70 protein and other antigens using linear regression and the Pearson correlation coefficient (r) (Fig. 5). There was a strong correlation (*P*<0.0001) between a level of Abs against V1V2_A244_-gp70 versus two other V1V2 fusion proteins (CaseA2 and ZM109) and three cyclic V2 peptides (A244, 92TH023, Du422), as well as gp120_A244_ and gp41_MN_. Thus, correlation of low level of Abs against V2 and gp120/ gp41 suggest that a deficiency of anti-V2 plasma Abs is related to a generally weaker Ab response to HIV-1 Env antigens in a subset of individuals infected with HIV-1.

**Figure 5.**
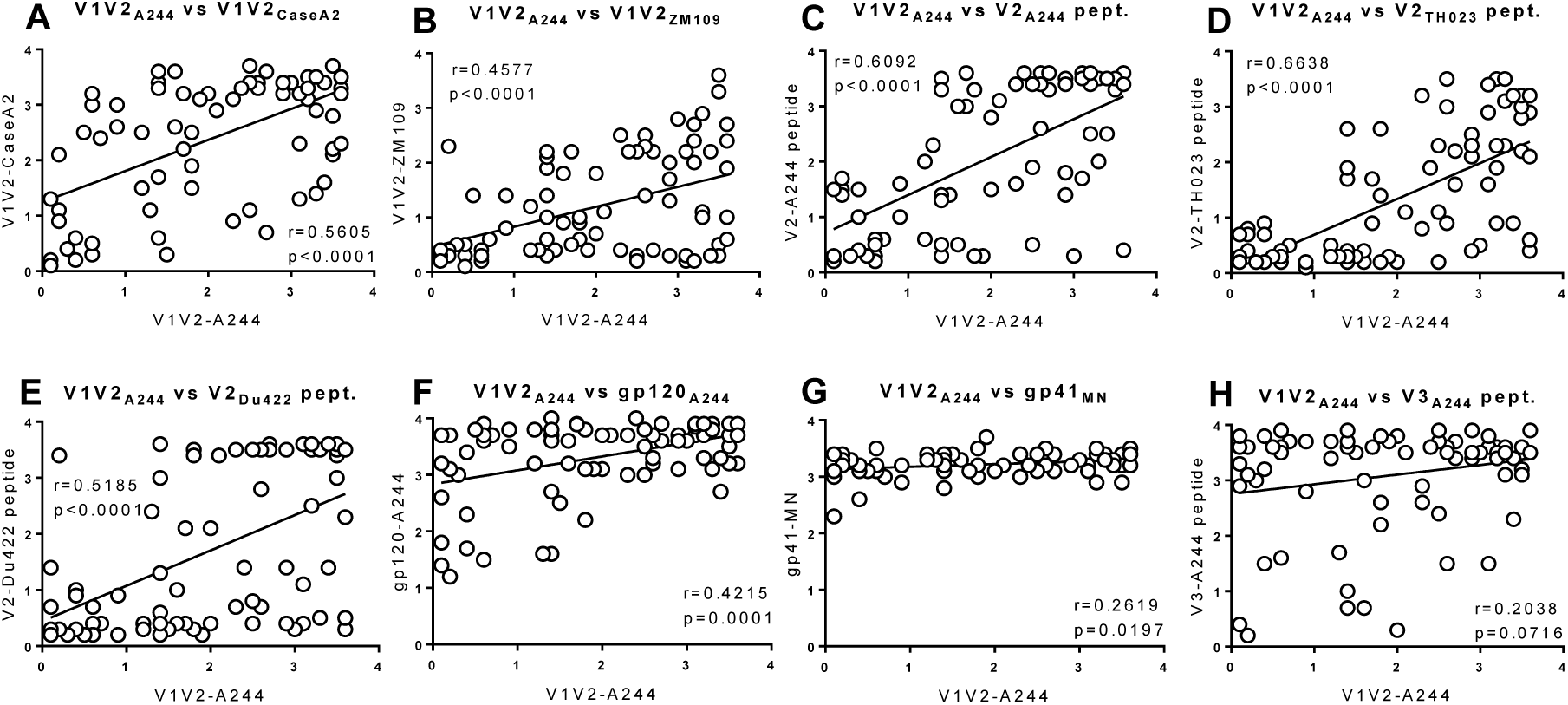
Correlation between binding activities (OD) of plasma Abs against V1V2_A244_-gp70 (V1V2_A244_) and other Env antigens. Plasma binding activity to V1V2_A244_ was correlated with: A) V1V2_CaseA2_-gp70; B) V1V2_ZM109_-1FD6; C) V2_A244_ peptide; D) V2_TH023_ peptide; E) V2_Du422_peptide; F) gp120_A244_; G) gp41_MN_, but not with H) V3_A244_ peptide. Correlation was determined by Pearson correlation coefficient (r) and by linear regression with P values.

### Characteristics of the V2 and V1 regions

We sequenced gp120 of the viruses in plasma of 12 selected patients to analyze the V2 and V1 regions, including properties that may have some influence on the immunogenicity of V2. From each plasma virus, 3 to 9 gp120 sequences were produced and used to generate consensus sequences for all 12 subjects (Table 2S). We analyzed the amino acid (AA) length, number of putative N-linked glycosylation sites, isoelectric point (pI), and charge of the V2 and V1 regions. For each region, we compared these features in the V2-deficient vs V2-reactive groups (Table 2, Table S3). In the V2 region, AA length was significantly longer in the V2-deficient vs V2-reactive group (43.5 vs 39.5 AA; *P*=0.0338), while pI (8.47 vs 9.25; *P*=0.1075) and charge (1 vs. 2.5; *P*=0.1613) were lower, but not significantly different in the V2-deficient vs the V2-reactive group, respectively (Table 3). In contrast to these data, the V1 region displayed statistically comparable AA length of 23.5 vs 24.5 (*P*=0.572), pI values of 4.89 vs 4.37 (*P*=0.6991) and charge: 1 vs –1.5 (*P*=0.8068) (Table 3).

**TABLE 3.** Characteristics of the V2 and V1 regions in HIV-infected individual who were had V2-deficient antibody responses or cross-reactive antibody response

| Features | V2 region |  | P<sup>1</sup> | V1 region |  | P<sup>1</sup> |
| --- | --- | --- | --- | --- | --- | --- |
|  | V2 Deficient Group | V2 Cross Group |  | V2 Deficient Group | V2 Cross Group |  |
|  | Median | (min, max) |  | Median | (min, max) |  |
| # amino acids (AA) | 43.5 (40, 58) | 39.5 (37, 41) | **0.034** | 23.5 (22, 32) | 24.5 (21, 38) | 0.572 |
| # glycosylation sites | 2 (2, 7) | 2 (0, 2) | 0.115 | 3 (2, 4) | 2.5 (1, 4) | 0.452 |
| pI | 8.47 (4.79, 9.66) | 9.25 (6.52, 9.69) | 0.108 | 4.89 (3.88, 9.37) | 4.37 (3.99, 9.58) | 0.699 |
| Charge | 1 (-1, 3) | 2.5 (0, 3) | 0.161 | -1 (-3, 2) | -1.5 (-4, 2) | 0.807 |
| α-helix K168-V172 | 1.78 (1.6, 1.9) | 1.57 (1.13, 2) | 0.376 | nt | nt | nt |
| β-sheet E153-I184 | -19.82 (-22, -16.3) | -20.32 (-22.97, -18.88) | 0.699 | nt | nt | nt |
| **Composite Indexes\*** |  |  |  |  |  |  |
| 1. AA-pI | 35.37 (31.98, 49.5) | 30.89 (27.87, 31.75) | **0.005** | 18.07 (13.63, 26.4) | 18.74 (14.42, 34.0) | 1.000 |
| 2. AA-Charge | 42 (40, 56) | 37.5 (35, 39) | **0.005** | 24.5 (21, 33) | 25.5 (22, 42) | 0.748 |
| 3. AA-pI+ Glyc. | 37.37 (33.98, 56.5) | 32.39 (27.87, 33.75) | **0.005** | 20.44 (16.63, 30.4) | 21.74 (15.42, 38) | 1.000 |
| 4. AA-pI- Glyc. | 33.37 (29.98, 42.5) | 29.39 (26.31, 29.75) | **0.005** | 16.07 (10.63, 22.4) | 15.74 (13.42, 30) | 1.000 |
| 5. AA-Charge+Glyc. | 44 (42, 63) | 39 (35, 41) | **0.005** | 27 (24, 37) | 28.5 (23, 46) | 0.936 |
| 6. AA-Charge-Glyc. | 40 (38, 49) | 36 (33, 37) | **0.005** | 22.5 (18, 29) | 22.5 (20, 38) | 0.808 |
| 7. AA-pI+helix | 37.2 (33.76, 51.4) | 32.39 (29.44, 33.7) | **0.002** | nt | nt | nt |
| 8. AA-Charge+helix | 43.82 (41.6, 57.9) | 38.99 (36.57, 40.13) | **0.002** | nt | nt | nt |

<sup>1</sup> P value, V2 deficient versus cross-reactive group (non-paired tests), Wilcoxon test; nt – not tested.
\*Composite indexes are based on difference or sum or linear combinations of more than one features; 1. # AA-pI; 2. # AA-Charge; 3. # AA-pI+ # glycosylation sites; 4. # AA-pI - # glycosylation sites; 5. # AA-Charge+# glycosylation sites; 6. # AA-Charge-# glycosylation sites; 7. # AA-pI+ α-helix propensity; 8. # AA-Charge+ α-helix propensity.

We also analyzed the β-sheet propensity for the central conserved segment (E153 to I184) and the α-helix propensity (K168 to V172) of the V2 region. The β-sheet propensity correlates with neutralization sensitivity of the virus (13) and may suggest a different immunogenicity profile, i.e., preferential presentation of different epitopes. The α-helix propensity is an estimate of the helical conformation, which is recognized by anti-V2 peptide mAbs such as CH58 (14) and putatively induces anti-V2p Abs; lower propensity suggest a higher tendency to form a helical structure. The two parameters were comparable between the V2-deficient and V2-reactive groups (Table 3, Table S3). Thus, the V2 consensus sequences from the V2-deficient compared with the V2-reactive group were slightly longer, with one additional N-linked glycosylation site on average and a tendency to a lower pI and electrostatic charge in the V2-deficient group.

### Multivariable analysis

Given that the majority of features characterizing the V2 region, except for length, are not significantly different between the V2-deficient vs V2-reactive groups, we tested whether the combination of multiple features could distinguish these two groups (Table 3). We defined composite indexes by combining 2 or 3 features to determine if the difference between the V2-deficient and V2-reactive groups, in the V2 and V1 regions, is significant, using the nonparametric Wilcoxon test. The difference between “No. AA” and “pI” and the difference between “No. AA” and “Charge” completely separated the V2-deficient from V2-reactive groups. That is, for each of the composite indexes, the maximum value (Max) in the V2-reactive group was lower than the minimum value (Min) in the V2-deficient group. Despite the small sample sizes, the lack of overlap in the values of each composite index between the two groups is statistically significant (*P*=0.005) (Table 3). Similarly, combining three features (“the sum of No.AA and No. glycosylation sites minus pI,” “the sum of No. AA and No. glycosylation sites minus charge,” and “the sum of No. AA and α-helix minus pI or charge” also separated the V2-deficient and V2-reactive groups; the non-overlapping index values between the groups are statistically significant (*P*=0.005 and 0.002, respectively (Table 3). This analysis provides evidence that multivariate predictions are more powerful than any single V2 variable for distinguishing V2-deficient from V2-reactive patient groups (Table 3). The same method was used for analysis of the V1 region as a control and did not show any significant differences between the two groups (Table 3).

## DISCUSSION

The present study analyzed Abs against the V2 region in terms of their frequency and level variation in plasma samples derived from 79 Cameroonian volunteers chronically infected with HIV-1. Given that some samples were negative for anti-V2 Abs, we focused on characterization of the V2 region of these individuals’ viruses to look for characteristics that might explain the deficiency of anti-V2 response. This analysis was prompted by the observation that levels of anti-V2 Abs were inversely correlated with the reduced HIV infection rate in recipients of the vaccine used in the RV144 clinical trial (1, 2) although levels of anti-V2 Abs measured by OD were not unusually high (9).

In the present study, the frequency of plasma anti-V2 Abs was between 53% and 85% for the three V1V2 fusion proteins and was comparable to results in the RV144 study, which reported 84% frequency, possibly due to using the same V1V2 fusion proteins for screening (9). In previous studies of sera of individuals chronically infected with HIV-1, anti-V2 Abs were detected at a much lower frequency (from 12% to 48%), but either the V2 antigens tested were different or plasma Abs were analyzed by different methods (4–7).

The level of anti-V2 Abs showed broad variations, in contrast to a relatively high level of Abs against V3, gp120, and gp41, as measured by OD and also by 50% titers (Fig. 1). Eight out of 79 (10%) plasma samples had no reactivity to any of the six V2 antigens in longitudinal studies over 8 to 54 months, while all plasma samples reacted to gp120 and gp41 (Fig. 2C). There was a strong correlation between ODs of plasma Abs binding to V2 antigens and to gp120/gp41 (Fig. 5). For example, plasma samples with high or low OD for gp120 and gp41 Abs also had high or low levels of anti-V2 Abs, respectively, suggesting that a patient’s capacity to mount Ab responses to HIV-1, in general, is associated with the level of anti-V2 Abs.

We hypothesized that, in addition to a low Ab response to Env proteins, the lack of anti-V2 Abs in several patients may depend on unusual characteristics of the V2 regions. Analysis of V2 sequences revealed that the V2 region of viruses from donors with V2-deficient versus V2-reactive Abs was significantly longer (*P*=0.0303), with 1 additional N-linked glycosylation site, on average. Usually, a longer V2 region is accompanied by an increased number of N-linked glycosylation sites (15–17). There is a correlation between the length and number of N-linked glycosylation sites, as has been observed in 41 pseudo-typed viruses in which neutralization-sensitive viruses to anti-V2 mAbs displayed shorter length and fewer glycosylation sites (18). Given that additional glycosylation sites in the V2 region confer resistance to neutralization by V2 mAbs, this may also have an impact on immunogenicity, resulting in weaker or lack of Ab response, particularly in subjects with poor humoral immune response to infecting virus.

The charge and pI of the V2 region were also lower in the V2-deficient versus V2-reactive panel of sequences (Table 3). Both these factors play also a role in anti-V3 mAbs; the pI of the VH CDR3 was significantly lower for human anti-V3 mAbs neutralizing Tier 2 and 3 pseudo-typed viruses vs non-neutralizing mAbs (19). For example, JR-FL.JB virus (Tier 2) was neutralized by 12 anti-V3 mAbs, and the pI of their VH CDR3 was significantly lower compared with that of 36 non-neutralizing V3 mAbs (data not shown); all these V3 mAbs were developed in our laboratory (19). Although the results do not correspond to the phenomenon of failed Ab response to the V2 region, they underline a functional role of electrostatic interactions between Ab and virus and, possibly, in immunologic recognition as well.

All 4 features—length, number of N-glycosylation sites, pI, and charge—were also analyzed in the V1 region for control; there were no significant differences or tendencies in either higher or lower values between the groups with V2-deficient and V2-reactive Abs (Table 2).

One type of anti-V2 Abs binds to V2 linear epitopes, which are detected using V2 peptides. These epitopes were defined by two human mAbs, CH58 and CH59, which were generated from recipients of the vaccine used in the RV144 clinical trial (14). Both mAbs bind to the helical structure of the V2 region, based on crystallographic analysis (14). The higher propensity to helical conformation in the central V2 segment (K168 to V172) in the V2-deficient versus V2-reactive group, which indicates higher flexibility, may be less immunogenic (13).

For most features, changes in the V2 region were minimal for the V2-deficient and V2-reactive groups; although length was significantly longer in the V2-deficient group. However, multivariable analysis including combination of 2 or 3 features, for example, differences in length and pI or charge and linear combinations of length, pI, and glycosylation sites, completely separated the two groups and resulted in a highly significant difference between V2-deficient and V2-reactive groups (Table 3). Thus, we conclude that the combination of these small differences in multiple features may be responsible for the lower immunogenicity of the V2 region resulting in lack of anti-V2 Abs in patients with significantly lower Ab response to the Env proteins.

In summary, our results show a broad range of binding activity of plasma Abs against 3 V1V2 fusion proteins and 5 cyclic V2 peptides, with a frequency of 53% to 85%, whereas in all plasma samples, Abs bound to heterologous gp120 and gp41. A strong correlation between anti-V2 and anti-gp120/ gp41 plasma Abs suggests that lack of V2 Abs depends in part on the general ability of patients to mount Ab responses to HIV-1 proteins. Sequence analysis of the V2 region from plasma viruses in the V2-deficient Ab group indicates that HIV vaccine immunogens containing V2 segments with fewer glycosylation sites and higher electrostatic charge may be advantageous in inducing higher titers of anti-V2 Abs.

## MATERIALS AND METHODS

### Ethics Statement

Blood samples from men and women infected with HIV-1 were received from the Medical Diagnostic Center (MDC), Yaoundé, Cameroon. Written informed consent forms were signed by all participants in the study and approved by the National Ethical Review Board in Cameroon. The study has been reviewed and approved by the Institutional Review Board of New York University School of Medicine, New York, USA.

### Specimens

The panel of 79 plasma samples from Cameroonian patients chronically infected with HIV-1 was randomly selected and received from the collaborative cohort shared between New York University School of Medicine, New York, NY, and the MDC, Yaoundé, Cameroon. Plasma samples were collected longitudinally over a period of 8 to 60 months and analyzed serologically. The donors were presumably infected with non-clade-B viruses, which were confirmed in 12 study subjects by sequencing their plasma viruses (Table 1). Participants enrolled in this study had been referred to the MDC in Cameroon for medical consultation due receiving an HIV-positive diagnosis. Acquisition of HIV-1 infection was self-reported as caused by heterosexual transmission in more than 95% of cases.

### Recombinant proteins and peptides

Seven recombinant proteins—V1V2_A244_-gp70 (clade AE), V1V2_case A2_-gp70 (clade B), V1ΔV2_A244_-gp70 (V2 deleted), V1ΔV2_case A2_-gp70 (V2 deleted), gp120_A244_, gp41_MN_ (clade B), and p24_HXB2_ (clade B) —were purchased from Immune Technology Corp., New York, NY. Biotinylated cyclic peptides —V2_A244_, V2_92TH023_ (CRF01_AE), V2_Du422_ (clade C), V2_230_ (CRF02_AG, plasma virus from donor MDC230, Table S2) and V3_A244_ —were purchased from Biopeptide Co., Inc., San Diego, CA. One biotinylated cyclic peptide, V2_200_ (CRF02_AG), representing the V2 sequence of plasma viruses from donor MDC200 (Table S2), was purchased from Science Exchange, Palo Alto, CA. V1V2_ZM109_-1FD6 (clade C) was produced as previously described (20).

### Antibody binding assay (ELISA)

The plasma samples were screened by standard ELISA against recombinant V1V2 fusion proteins, gp120, gp41 and p24 and against biotinylated V2 and V3 peptides according to the methods in (21, 22). In short, proteins were coated directly onto Immulon 2Hb plastic plates at a concentration of 1 μg/mL. After overnight incubation at 4 ^°^ C, the plates were washed with 1X phosphate-buffered saline (PBS) containing 0.05% Tween 20 and blocked with PBS containing 2.5% bovine serum albumin and 7.5% fetal bovine serum. Plasma at dilution 1:100 was incubated with antigens in the plate for 1.5 h at 37^°^C. The plates were washed and incubated with alkaline phosphatase-conjugated goat anti-human IgG (γ specific) (Southern Biotech, Birmingham, AL) followed by washing and incubation with substrate to develop color, and read at 405 nm. IgG subclasses and IgA of plasma Abs against V1V2 fusion protein were detected using mouse mAbs against IgG subclasses and goat anti-human IgA (Southern Biotech). For screenings of plasma against biotinylated peptides, the peptides were immobilized at a concentration of 1 μg/mL onto streptavidin-coated plates (StreptaWell plates, Roche). The 50% titers of plasma Abs were determined by measuring the dilutions of plasma required for 50% maximal binding by linear regression (23). A positive reaction by a plasma sample at 1:100 dilution was defined as an OD of the mean + 3 standard deviations from the 6 plasma samples of healthy individuals from Cameroon, and then rounded to 1 tens digit.

### CD4^+^ T cell counts

The CD4^+^ T cell counts were determined using the Guava EasyCD4 Test (Guava Technologies Inc, Hayward, CA) with 2-color direct and absolute counting of number cells per μL (24).

### Viral load

Viral load was determined using Abbott m2000 Real Time HIV-1 assays according to the manufacturer’s instructions (Abbott Molecular, Des Plaines, IL).

### Sequencing of the virus envelope

Viruses in 500 μL plasma samples were concentrated by centrifugation for 2 h, 14,000 rpm, at 15°C prior to RNA extraction. After removal of 360 μL supernatant, the virus pellet was resuspended in the remaining 140 μL supernatant by vortexing, and viral RNA was extracted using the QIAamp viral RNA mini kit (Qiagen Inc., Valencia, CA) as per manufacturer’s instructions. Reverse transcription and nested PCR were performed using the SuperScript One-Step RT-PCR system with platinum *Taq* and Platinum PCR high-fidelity SuperMix (Invitrogen, Carlsbad, CA), respectively, as per manufacturer’s instructions, to isolate a portion of *env* (gp120 + fragment of gp41), HXB2 region 6225-7817 (∽1600 bp). PCR products were cloned into the pCR4 TOPO cloning vector (Life Technologies, Carlsbad, CA) and transformed into One Shot TOP10 competent *E. coli*. Positive clones were cultured in LB_Amp_ medium, and plasmids were isolated using the QIAprep Spin Miniprep kit (Qiagen Inc). Plasmids were sequenced for the insert portion using universal primers M13F/R (pCR4 TOPO). Sequence analysis and alignment was performed using DNASTAR and Clustal W (25). Consensus sequences were generated using the Los Alamos National Library program ConsensusMaker.

### Statistical analysis

The nonparametric Mann-Whitney-Wilcoxon test was used for comparing OD, titers of plasma Abs, and composite indexes based on linear combinations of individual characteristics or features (Table 3). Each individual feature or characteristic of the V2 region (e.g., isoelectric point, charge, number of glycosylation sites) has limited power to distinguish the V2-deficient group from the V2-reactive group. However, composite indexes based on simple difference or sum or linear combinations of more than 1 feature (e.g., No_AA-pI in Table 3) can separate these groups. The relationship of plasma Ab binding to different antigens was determined by Pearson correlation coefficient (r) with *P* values and by linear regression. Statistical analysis and graphing of the data were generated using GraphPad Prism version 7 (GraphPad Software, La Jolla, CA).

## ACKNOWLEDGMENTS

The study was supported by NIH grants AI112546 (MKG) and AI100151 (SZP/XPK). The authors thank the individuals with HIV-1 infection for their donation of blood samples for this study, which were collected at the Medical Diagnostic Center, Yaoundé, Cameroon. We would like to thank Dr. Michelle Ryndak for reviewing the manuscript. L.L., L.L., A.N., L.M.M., S.S., A.K., L.A., X-H.W., M.T. and Y.S. performed research; Y.S., M.T., S.Z-P., X-P.K., R.D. and M.K.G. analyzed data; and M.K.G. wrote the paper.

